# Missense variants in the myosin binding domains of *MYBPC3* and *MYBPHL* impair sarcomere incorporation

**DOI:** 10.1101/2025.09.29.679322

**Authors:** Kelly N. Araujo, Hannah E. Cizauskas, Yoldas Yildiz, Geena E. Fritzmann, Tuan Bui, Lucas M. Wittenkeller, Alexandra Peña, Toni R. Pak, David Y. Barefield

## Abstract

Approximately 40% of genetic hypertrophic cardiomyopathy cases involve mutations in *MYBPC3*, which encodes cardiac myosin binding protein-C (cMyBP-C), a key regulator of sarcomere contractility. The atrial-specific paralog, myosin binding protein-H like (MyBP-HL), has been associated with dilated cardiomyopathy in humans and mice. Both proteins bind to the same binding sites in the thick filament C-zone. In the atria, cMyBP-C and MyBP-HL are found at ∼1:1 ratios, while ventricles only express cMyBP-C, which is found at twice the atrial level, indicating a stoichiometric relationship. In the atria, we hypothesize that missense variants in either gene may cause alterations in thick filament binding affinity and changes in the normal ∼1:1 ratio. Notably, MyBP-HL deletion in atrial myofibrils accelerates relaxation kinetics, suggesting that altered stoichiometry impacts biophysical parameters.

We hypothesized that deletion, overexpression, or missense variants in either gene would alter the abundance of the other protein in atrial sarcomeres, affecting sarcomere localization and function. To test this, we engineered two constructs: a mini-C construct comprising thick filament-binding domains of cMyBP-C and a MyBP-HL construct. Selected *MYBPC3* and *MYBPHL* missense variants were introduced and expressed in neonatal rat ventricular cardiomyocytes (NRVMs). Sarcomere localization was assessed by co-localization with endogenous cMyBP-C. *MYBPC3* variants were selected across a range of pathogenicity, while MYBPHL variants were based on evolutionary conservation of residues. *MYBPHL* variants Gly275Ser, Arg285His, and Ala342Thr induced significant sarcomere mislocalization, and *MYBPC3* variants Pro1181Ala and Asn1257Lys showed variable effects on sarcomere mislocalization.

To assess stoichiometric effects, we developed a T2A/P2A polycistronic construct to co-express mini-C, Td-Tomato, and MyBP-HL. Immunoblotting and mass spectrometry confirmed consistent and reproducible expression. We identified several *MYBPC3* and MYBPHL variants that reduced the affinity of their protein for myofilament incorporation. These results suggest that this 2A construct is a useful tool for measuring the effect of myosin binding protein missense variants on sarcomere affinity, with implications for assessing pathogenicity of these variants.

## Introduction

Cardiomyopathy is a common heart disease that often leads to heart failure and subsequent mortality [1]. Mutations in sarcomere genes play a major role in both hypertrophic cardiomyopathy (HCM) and dilated cardiomyopathy (DCM) [2-4]. HCM primarily arises from alterations in sarcomere proteins that lead to hypercontractile, thickened, and stiff ventricles [3, 5] while DCM involves mutations in many cellular mechanisms that result in ventricular wall thinning, left ventricular dilation and impaired contractility [6-8]. HCM and DCM frequently coexist with atrial fibrillation, a comorbidity that exacerbates disease progression [9-11]. Atrial fibrillation has a prevalence of 20 – 30% in patients with HCM and DCM, 4 – 6 times more frequently than in the general population [12-15]. While cardiomyopathy-causing mutations can affect atrial function, their specific impact on atrial contractility and the pathogenesis of atrial fibrillation remains poorly understood [16]. This is especially the case in genes that encode chamber-specific or chamber-enriched protein isoforms.

Recent studies have shown that the proteins encoded by the well-characterized HCM-associated gene, *MYBPC3*, and the atrial-enriched, DCM-linked gene, *MYBPHL*, directly compete for incorporation into the atrial sarcomere [17]. The sarcomere is comprised of a dozen core proteins that are organized in a semi-crystalline structure with a defined stoichiometry [18]. Myosin binding protein H-like (MyBP-HL) is highly homologous to cardiac myosin binding protein-C (cMyBP-C) via their thick filament binding domains that allow their proper sarcomere incorporation [17]. In atrial sarcomeres, an increase or decrease in expression of either MyBP-HL or cMyBP-C causes a respective decrease or increase in the other. Nonsense mutations in *MYBPC3* and *MYBPHL* both prevent proper sarcomere localization [17, 19]. In contrast, missense mutations in *MYBPC3* have variable pathogenic effects depending on the location of the mutation within the protein, with some mutations within the thick filament binding domains causing improper sarcomere incorporation [19]. However, thick filament disrupting myosin binding missense mutations have only been studied in the context of the ventricle, where *MYBPC3* is the only myosin binding protein gene expressed.

We aimed to determine whether the presence of missense variants within the thick filament binding domains of either MyBP-HL or cMyBP-C will alter the abundance of the other myosin binding protein within atrial sarcomeres. To investigate this, we have selected *MYBPC3* and *MYBPHL* variants from the GnomAD database and screened their effects on myosin binding protein localization within the sarcomere. To study the effects of myosin binding protein missense variants on sarcomere incorporation, we developed a 2A self-cleaving peptide construct to simultaneously express both myosin binding proteins. We use this tool to test *MYBPC3* or *MYBPHL* missense variants and determine changes in myosin binding protein stoichiometry.

## Materials and Methods

### NRAM/NRVM isolation

All animals were housed and sacrificed in accordance with Loyola University Chicago’s Institutional Animal Care and Use Committee in adherence to the US National Institutes of Health Guide for Care and Use of Laboratory Animals. Neonatal rat hearts were isolated from 0 – 1-day old Sprague-Dawley pups (Charles River Laboratories). Atria and ventricles were separated, minced, and digested at 37 °C in Krebs-Henseleit buffer with collagenase and 0.05% trypsin with occasional swirling for 15 minutes. This process was repeated 5 times for atria and 6 times for ventricles. At each time point, the buffer was collected, strained, and added to a separate tube containing neonatal rat isolation media (Dulbecco’s Modified Eagle’s Medium (DMEM, Gibco) supplemented with 10% Fetal Bovine Serum (FBS) and 1% penicillin-streptomycin) to stop the digestion. Digested cardiomyocyte were then centrifuged at 800 × g for 10 minutes and supernatant was aspirated. Cells were resuspended in 10 mL of neonatal rat cardiomyocytes isolation media (high glucose DMEM, 10% FBS, 1% penicillin-streptomycin). The cell suspension was then plated and incubated at 37 °C for 1.5 hours to separate the fibroblasts from cardiomyocytes. The media, containing the isolated cardiomyocytes, were then collected and centrifuged at 500 × g for 10 minutes. Supernatant was aspirated and the cell pellet was resuspended in neonatal rat cardiomyocyte maintenance media (DMEM/F12, 18.5% M199 medium, 5% horse serum, 1% FBS, 1% penicillin-streptomycin, 1% insulin-transferrin-selenium, 10 μM BrdU) in a total volume of 20 mL for neonatal rat ventricular cardiomyocytes (NRVMs) and 5 mL of neonatal rat atrial cardiomyocytes (NRAMs). Cells were then counted and brought to 2 million cells/mL for protein collection and 1 million cells/mL for adherence to coverslips.

### Plasmid design

We used the human *MYBPC3* sequence from domains C8 to C10 inserted into the pCMV6 expression vector to create the mini-C construct. For the MyBP-HL construct, the full-length human *MYBPHL* sequence was inserted into the pCMV6 expression vector. We used the information reported by the GnomAD database about human *MYBPC3* and *MYBPHL* missense variants and mimicked those chosen missense variants in mini-C and *MYBPHL* plasmid sequences, respectively. We designed specific primers for site-directed mutagenesis to generate missense variants in human *MYBPHL* (**Table 1**) and mini-C (**Table 2**) using Q5 Hot Start High-Fidelity DNA polymerase (New England Biolabs M0494S). Plasmids were sequenced by Sanger reaction (Azenta Life Sciences, Burlington, MA). Transfection quality plasmids were prepared using an endotoxin-free maxi prep kit (Qiagen, 12362).

**Table 1.**
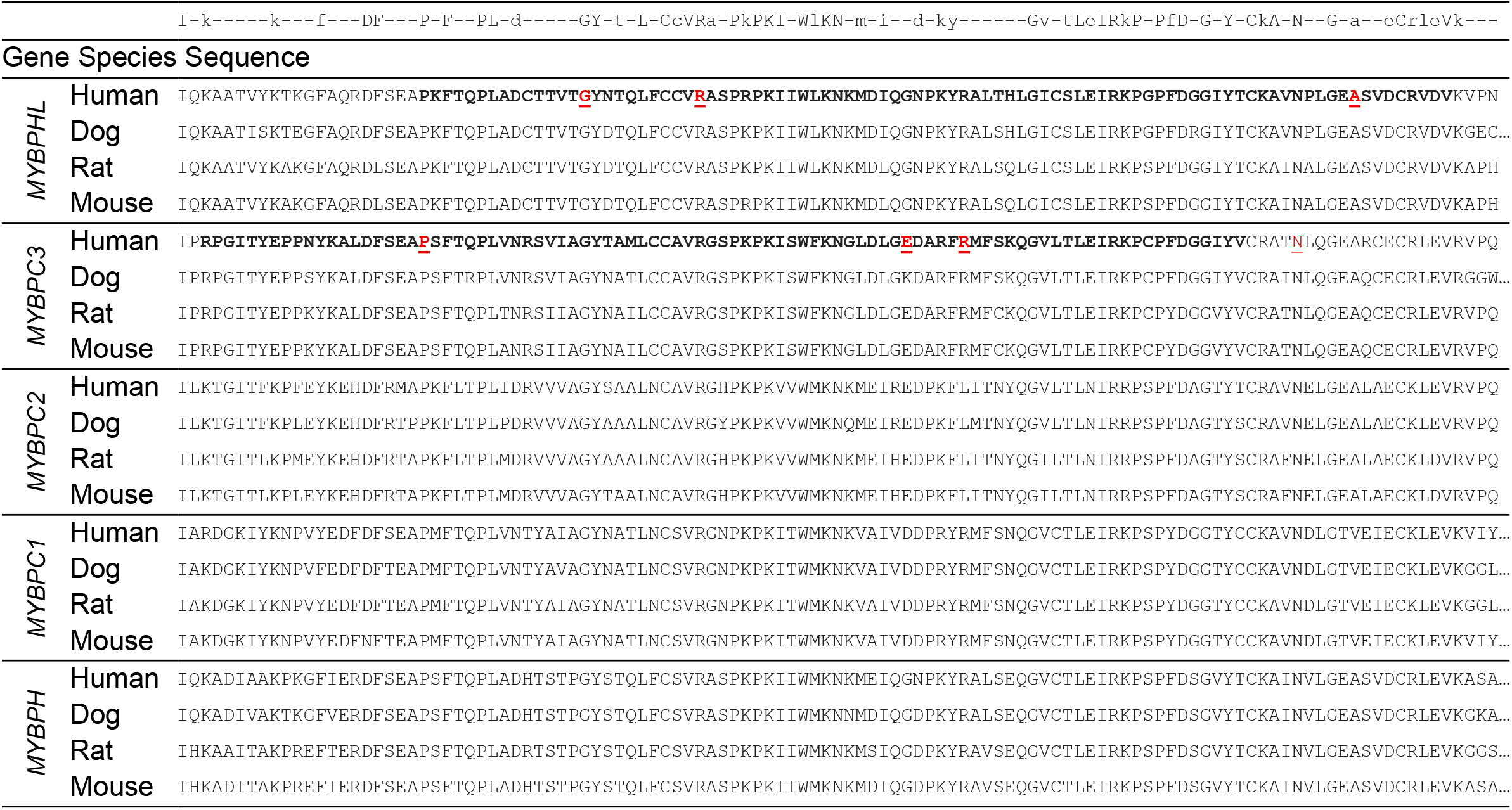
Consensus sequence of the penultimate immunoglobulin domain of the five myosin binding proteins across four species. Capital letters indicate amino acids that have complete or nearly complete conservation. Lower case letters indicate lower conservation with the most common residue marked. Bold lines for human *MYBPHL* and *MYBPC3* indicate residues that comprise the HL3 or C10 immunoglobulin domain, respectively. Red, underlined letters mark the location of the missense variants.

**Table 2.**
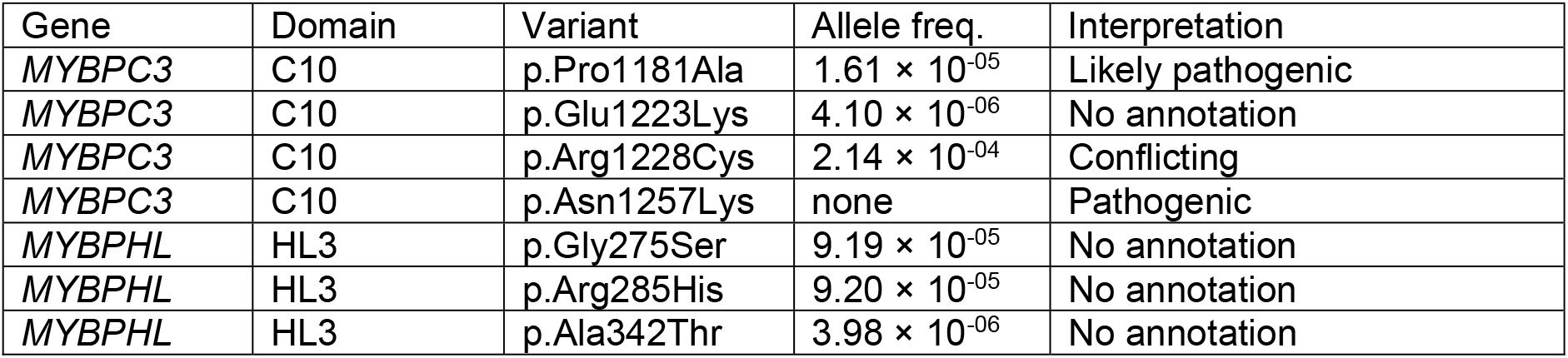
*MYBPC3 and MYBPHL va*riants with varying pathogenicity from GnomAD database.

| Gene | Domain | Variant | Allele freq. | Interpretation |
| --- | --- | --- | --- | --- |
| *MYBPC3* | C10 | p.Pro1181Ala | $1.61 \times 10^{-05}$ | Likely pathogenic |
| *MYBPC3* | C10 | p.Glu1223Lys | $4.10 \times 10^{-06}$ | No annotation |
| *MYBPC3* | C10 | p.Arg1228Cys | $2.14 \times 10^{-04}$ | Conflicting |
| *MYBPC3* | C10 | p.Asn1257Lys | none | Pathogenic |
| *MYBPHL* | HL3 | p.Gly275Ser | $9.19 \times 10^{-05}$ | No annotation |
| *MYBPHL* | HL3 | p.Arg285His | $9.20 \times 10^{-05}$ | No annotation |
| *MYBPHL* | HL3 | p.Ala342Thr | $3.98 \times 10^{-06}$ | No annotation |

**Table 3.**
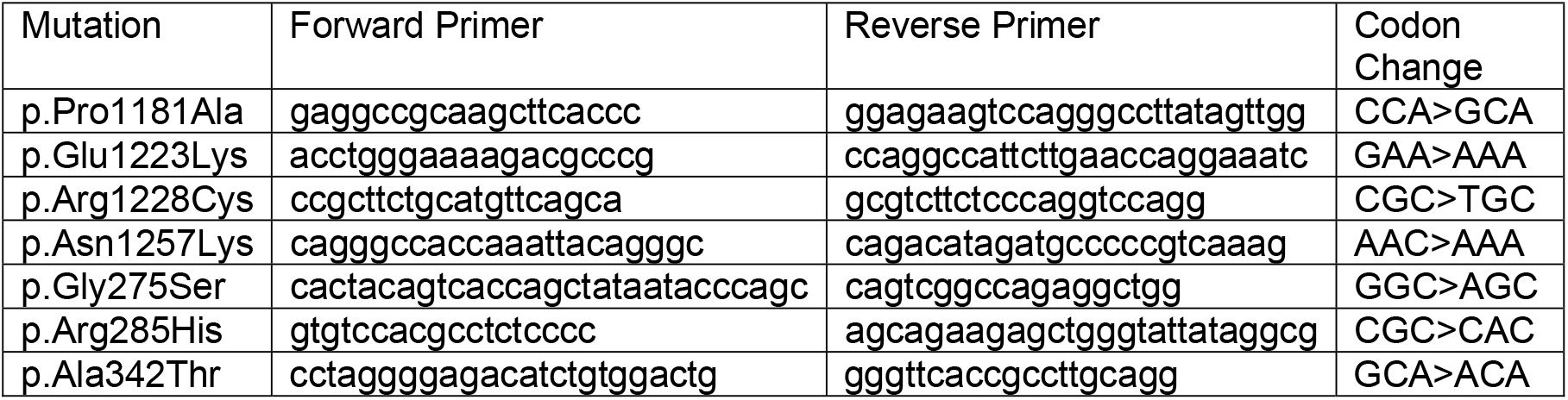
Site-directed mutagenesis primers used to introduce *MYBPC3* and *MYBPHL* missense variants within MyBP-HL and T2A/P2A constructs.

### Myofilament Fractionation

Neonatal rat ventricular cardiomyocytes were seeded at 2 × 10^6^ cells per well in 6-well cell culture plates. NRVMs were transfected with 3.14 μg of plasmid using Lipofectamine 3000 (ThermoFisher, L3000008). NRVMs were scraped off plates with F-60 buffer (60 mM KCl, 30 mM imidazole, 2 mM MgCl_2_) + 0.1% triton + protease inhibitor (ThermoFisher, A32955) and then spun down at 12,000 × g for 10 minutes and the supernatant was kept as the supernatant samples. This removed the soluble protein fraction, leaving the myofilament proteins in the pellet. The pellet was washed with F-60 buffer + protease inhibitor + 10% triton X-100 3 times. The pellet was then resuspended in 8 M Urea + 20 mM dithiothreitol + 0.2% SDS + protease inhibitor. To homogenate the myofilament proteins, the samples were placed in a water sonicator for 3 cycles of 30 seconds of on and off sonication. For normalization, 30 µL of myofilament samples were mixed with 10 µL of 4× dye. Supernatant samples were standardized using the BCA protein assay (ThermoFisher, 23227).

### Immunoblotting

Samples were resolved by 10% SDS-PAGE and transferred onto PVDF membranes (ThermoFisher, 88520). Membranes were blocked with tris-buffered saline with 1% casein blocker (Biorad, 610782) for one hour. Primary antibodies against human cMyBP-C (Santa Cruz, 137180), MyBP-HL (Barefield, 2022; 1:2500) and Anti-DDK (Origene, TA150078), phosphorylated serine 273 cMyBP-C [90], actinin (ProteinTech, 14221-1-AP) were incubated in blocking buffer overnight at room temperature. Secondary goat anti-rabbit (Jackson ImmunoResearch, 115-035-003) was probed in blocking buffer for one hour. Membranes were incubated with HRP substrate Clarity western ECL (Biorad, 1705062) for 10 seconds, and luminescence was captured with a BioRad ChemiDoc MP imaging system. After specific detection of proteins, blots were stained for total protein using Coomassie stain by incubation in fixating solution (50% methanol, 7% acetic acid, 43% water) for 5 – 10 minutes and then stained with Coomassie for 20 – 30 minutes. Blots were rinsed in water and to reduce background, blots were rinsed with destain solution (50% methanol, 40% water, 10% acetic acid) for 5 – 10 minutes each rinse and left overnight to dry and imaged the following day.

### Immunofluorescence staining and confocal microscopy

For sarcomere localization imaging, 1.75 × 10^5^ myocytes were plated on 12 mm glass coverslips (Fisher Scientific, 12-541-000) coated with GelTrex membrane matrix (ThermoFisher, A1413202). Cells were transfected with 1 μg of MyBP-HL construct, mini-C construct, MyBP-HL missense variants, or mini-C missense variants using Lipofectamine 3000 (ThermoFisher, L3000008). Cells were fixed with 4% paraformaldehyde (ThermoFisher, 50-980-487) and permeabilized with 0.25% Triton X-100 (Millipore Sigma, X100). Cells were blocked with 20% fetal bovine serum, 0.1% Triton in phosphate-buffered saline. Primary antibodies against MyBP-HL (Barefield et al., 2022; 1:2,500), Anti-DDK (Origene, TA150078) and cMyBP-C (Santa Cruz Tech, E7; sc-137180; 1:2500) were incubated in blocking buffer overnight at 4 °C. Secondary antibodies donkey anti-mouse IgG Alexa Fluor 488 (ThermoFisher, A21202; 1:2500) and donkey anti-rabbit IgG Alexa Fluor 568 (ThermoFisher, A10037; 1:2500) were incubated in blocking buffer for one hour at room temperature. Nuclear staining was done with Hoechst dye for 10 minutes. For confocal microscopy, coverslips were mounted using ProLong Gold (ThermoFisher, P36930). Images were obtained using a Zeiss Axio Observer microscope, connected to a W1 Confocal Spinning Disk system (Yokogawa) equipped with Mesa field flattening (3i), a motorized X,Y stage (ASI), a Prime 95B sCMOS camera (Photometrics), and a multifiber laser launch (3i). We used 488 nm and 561 nm lasers for confocal imaging of NRMs. Imaging was performed using either a 63×, 1.4 NA Plan Apochromat or a 10×, 0.45 NA Plan Apochromat objectives (Zeiss). The microscope operations were controlled through Slidebook 6 Software (3i). Images were processed using a Z-plane average picture of MyBP-HL and cMyBP-C channels. Then, cMyBP-C positive images were used to select sarcomere doublets, and we determined regions of interest. Colocalization analysis was performed using Coloc2 plugin for ImageJ. Data are expressed as a correlation between cMyBP-C and MyBP-HL channels using a Pearson’s R index correlation coefficient.

### Peptide preparation

NRVMs were seeded at 1.5 × 10^6^ cells per well in 6-well cell culture plates. NRVMs were transfected with 3.14 μg of T2A/P2A plasmid as described in previous section. NRVMs were dissociated using 0.25% trypsin and collected in NRM maintenance media. Cells were spun down at 2000 × g for 4 minutes, supernatant discarded, and cell pellets were washed with sterile cold phosphate-buffered saline, supernatant was discarded, and pellets were resuspended in 8 M urea in 50 mM ammonium bicarbonate (pH 7.8). Cell pellets were sonicated using Bioruptor Pico sonicator (Diagenode) and underwent 6 cycles of sonication, 15 seconds on and 30 seconds off, for 4 minutes and 30 seconds. For trypsin digestion, protein samples were vortexed for 1 hour (Eppendorf ThermoMixer F1.5). 5 mM dithiothreitol (Sigma-Aldrich) was added to sample and incubated at 37 °C for 1 hour (Eppendorf ThermoMixer F1.5) to reduce disulfide bonds. To alkylate the samples, the samples were then incubated in 15 mM iodoacetamide (Sigma-Aldrich) for 30 minutes at room temperature. To quench the excess iodoacetamide, 25 mM dithiothreitol was added to samples and incubated at room temperature for 15 minutes. Protein samples were then diluted 1:4 with 50 mM ammonium bicarbonate (pH 7.8) to reduce the urea concentration. Samples were then digested and incubated with mass spectrometry grade proteases (ThermoFisher) (1:50 wt/wt) overnight at 37 °C. Peptides were then purified and desalted using C18 spin columns (G-Biosciences), speed-vacuumed dried, and stored at −80 °C until further use. To determine peptide quantification, peptides were reconstituted in 0.1% formic acid and quantified (Thermo Scientific).

### Mass Spectrometry

1.0 µg of resolubilized peptides were loaded onto a Vanquish Neo ultra-high-performance liquid chromatography system (Thermo Fisher) with a heated trap and elute workflow (c18 PrepMap, 5 mm, trap column, G-Biosciences, 160454) in a forward-flush configuration connected to a 25 cm Easyspray analytical column (Thermo Fisher ES802A rev2) 2 µM,100 A, 75 µm x 25 µm with 100% buffer A (0.1% formic acid in water) and the column oven operating at 40 °C. Peptides were eluted over a 150 minute gradient using 80% acetonitrile, and 0.1% formic acid (buffer B), going from 4% to 15% over 10 minutes, to 40% over the next 90 minutes, then to 55% over the next 37 minutes, then to 99% over 6 minutes, and then kept at 99% for 7 minutes, after which all peptides were eluted. Spectra were acquired with an Orbitrap Eclipse Tribrid mass spectrometer (MS) with field asymmetric ion mobility spectrometry pro interface (ThermoFisher) running Tune 3.5 and Xcalibur 4.5. For all acquisition methods, spray voltage was set to 2000 V, and ion transfer tube temperature was set at 300 °C. Field asymmetric ion mobility spectrometry switched between compensation voltages of −45 V, – 55 V, and −65 V with cycle times of 1.5 seconds. MS1 spectra were acquired at 120,000 resolutions with a scan range from 375 to 1600 m/z, normalized AGC target of 300%, and maximum injection time of 50 ms, S-lens RF level set to 30, without source fragmentation and datatype positive and profile. Precursors were filtered using monoisotopic peak determination set to peptide; included charge states: 2 – 7 (reject unassigned); dynamic exclusion: enabled with n = 1 for 60 seconds exclusion duration at ten ppm for high and low. data dependent MS/MS scan using isolation mode: quadrupole; isolation window (m/z): 1.6; activation type: HCD with 30% collision energy; IonTrap detector with scan rate: turbo; AGC target: 10,000; maximum injection time: 35 ms, micro scans: 1; and data type: centroid.

Raw data were analyzed using Proteome Discoverer 2.5 (ThermoFisher) using Sequest HT search engines. The data were searched against the Homo sapiens Uniprot protein sequence database (UP000005640). The search parameters included precursor mass tolerance of 10 ppm and 0.06 Da for fragments, allowing two missed trypsin cleavages, oxidation (Met) and acetylation (protein N-term), and phosphorylation (S, T, Y) as variable modifications, and carbamidomethylation (C) as a static modification. Percolator PSM validation was used with the following parameters: strict false discover rate of 0.01; relaxed false discover rate of 0.05; maximum ΔCn of 0.05; and validation based on q-value. Precursor ions quantifier settings were to use unique + razor for peptides; precursor abundance was based on intensity, normalization based on total peptide amount, protein abundance was calculated by the summed intensity of connected peptides, and protein ratios were calculated based on protein abundance and a background-based T-test was used to calculate statistical significance.

## Statistical Analysis

All data were analyzed using Prism8 software (GraphPad, San Diego, CA). n = corresponds to number of different NRM isolations. We plotted an average of n ± SEM; an ordinary one-way ANOVA and subsequent Dunnett’s comparison test were performed. For cMyBP-C and MyBP-HL colocalization we used a n = 4; an ordinary one-way ANOVA and subsequent Dunnett’s multiple comparison test were performed. Densitometry analysis was performed using Fiji software, and data were analyzed with Prism8 with a n = 3; ordinary one-way ANOVA and subsequent Holm-Sídák multiple comparison test was performed.

## Results

To investigate whether missense mutations in the thick filament binding domains of MyBP-HL and cMyBP-C shift the myosin binding protein stoichiometry, we have selected different *MYBPC3* variants from the gnomAD genetic database with various levels of interpretations, ranging from no annotation to pathogenic. All mutations are located within the C10 domain of cMyBP-C (**Table 2**). We selected a positive control, *MYBPC3* Asn1257Lys, which was shown to cause improper sarcomere incorporation of cMyBP-C [19]. We also selected *MYBPHL* variants in highly conserved residues from gnomAD (**Table 1**). All mutations are localized within the HL3 domain, which corresponds to the C10 domain in *MYBPC3*. We hypothesize that the variants with high levels of pathogenicity or in highly conserved residues for cMyBP-C and MyBP-HL, respectively, would be the most likely to result in sarcomere mislocalization.

### Mini-C and MyBP-HL constructs co-localize with endogenous cMyBP-C within NRVMs

To investigate the effects of selected missense variants on cMyBP-C and MyBP-HL, we designed constructs capable of integrating into the sarcomere, allowing us to assess the effects of missense variant. While we could have used the full-length cMyBP-C sequence, we wanted to create a construct that would be the same size as MyBP-HL with only the thick filament binding domains to ensure that any changes in sarcomere localization would be due to the expression of missense variants and not due to size. Thus, we created a construct solely consisting of the thick filament binding domains of cMyBP-C with a DDK tag on its N-terminal end, named the “mini-C” construct. The MyBP-HL construct consists of the full-length human MyBP-HL with a DDK tag on its N-terminal end (**Fig. 1A**).

**Figure 1.**
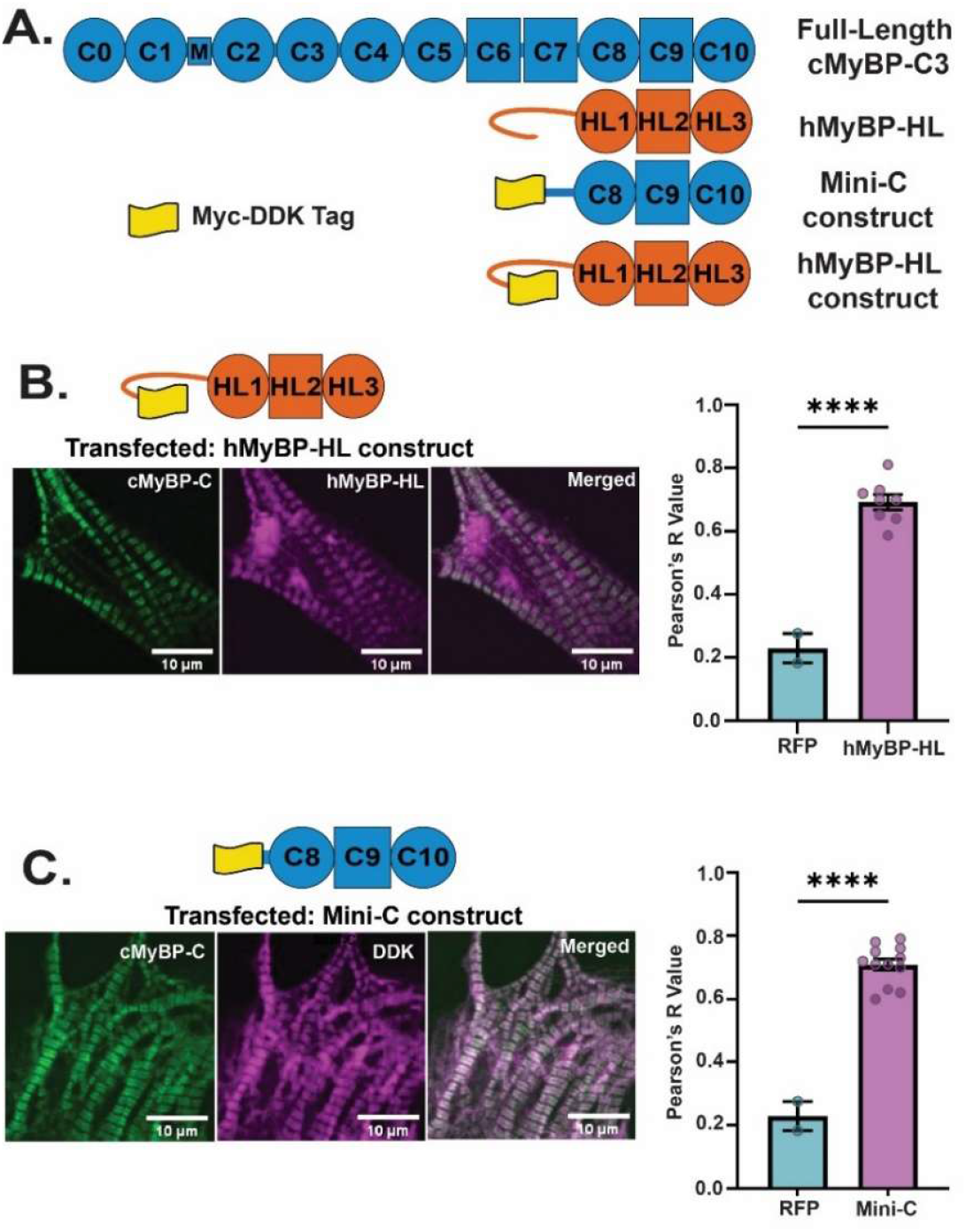
Mini-C and MyBP-HL properly incorporate into the sarcomere in NRVMs. (A) Domain schematic showing full-length cMyBP-C, full-length MyBP-HL, mini-C construct and MyBP-HL constructs. The mini-C construct consists of the three C-terminal domains found in full-length cMyBP-C, critical for myosin binding, and a Myc-DDK tag on the N-terminus. The MyBP-HL construct consists of the full-length MyBP-HL with a Myc-DDK tag on the N-terminus. (B) MyBP-HL construct transfected in neonatal rat ventricular cardiomyocytes and proper sarcomere localization was determined via immunofluorescence staining for endogenous cMyBP-C and MyBP-HL. Co-localization was quantified via Pearson’s R value (N = 2; One-way ANOVA). (C) Mini-C construct transfected in NRVMs and stained using Anti-DDK. Proper sarcomere localization was determined and quantified as previously described in (B). (N = 4; One-way ANOVA).

To ensure that the constructs properly incorporate into the sarcomere, we transfected NRVMs and compared the construct localization with endogenous cMyBP-C via immunostaining. Both mini-C and MyBP-HL constructs were shown to co-localize to the C-zone by their clear overlap with endogenous cMyBP-C (**Fig. 1B**). Co-localization was quantified by Pearson’s R-value, measuring the strength and direction of a linear relationship between the spatial correlation of endogenous cMyBP-C and the constructs. Both mini-C and MyBP-HL constructs show high Pearson’s R-value, indicative of a high co-localization with endogenous cMyBP-C (**Fig. 1C**).

### *MYBPC3* missense variants expressed in mini-C construct cause various levels of sarcomere mislocalization

To evaluate the effect of missense variants on sarcomere localization, we transfected NRVMs with wild-type mini-C and mini-C expressing the selected *MYBPC3* missense variants (**Fig. 2A**). Sarcomere localization was evaluated by immunofluorescence microscopy using endogenous cMyBP-C to mark the C-zone and co-localization was quantified using Pearson’s R Value (**Fig. 2B, C**). The Glu1223Lys and Arg1228Cys variants showed no changes in sarcomere localization. Variant Pro1181Ala caused some improper localization but with high variability that did not reach statistical significance. Lastly, the positive control Asn1257Lys caused complete mislocalization from the sarcomere, aligning with its pathogenic annotation.

**Figure 2.**
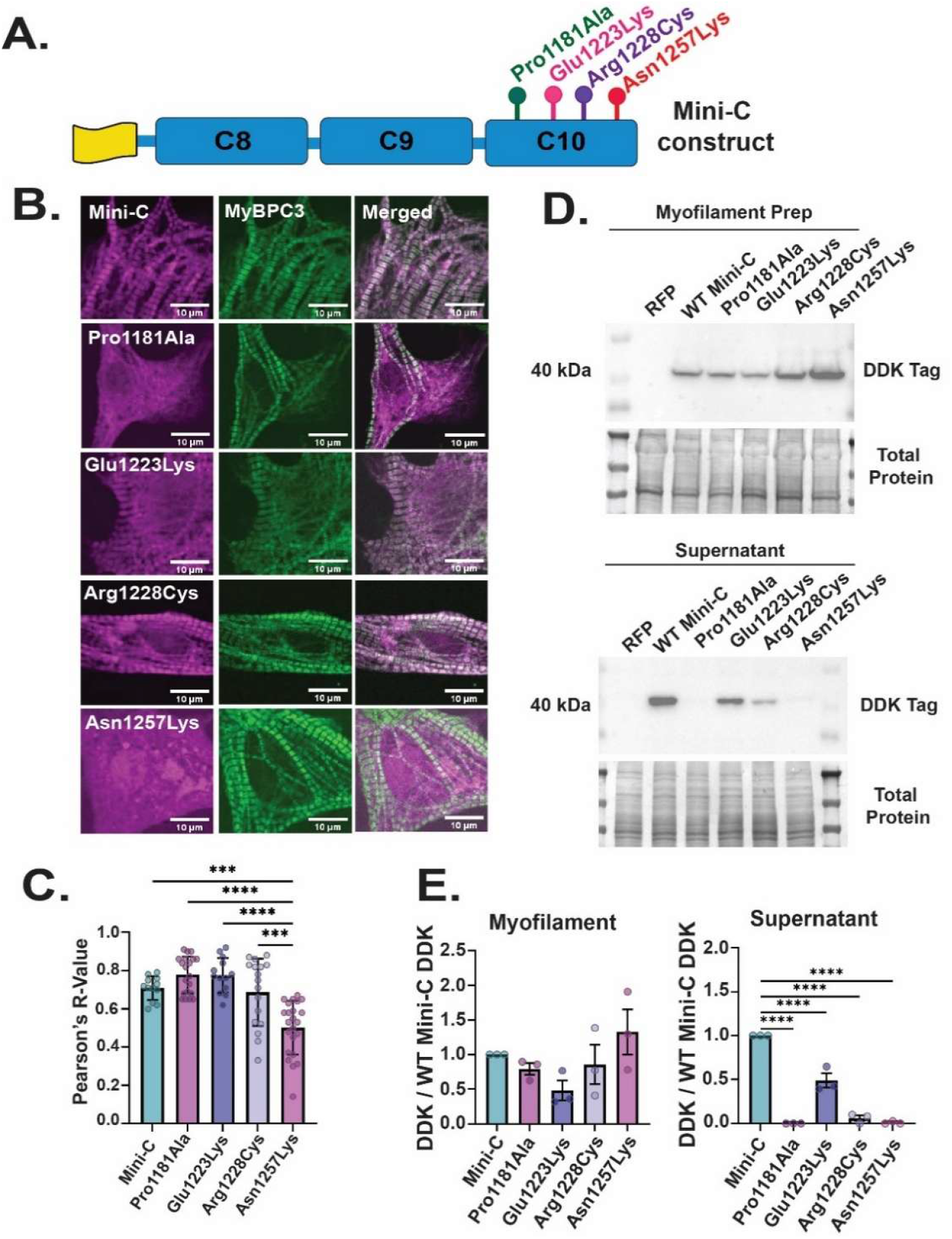
Missense variants within Mini-C impair sarcomere localization within NRVMs. (A) Schematic showing presence of selected missense variants within mini-C construct, all located in the C10 domain. (B) NRVMs were transfected with the missense variants expressed within the Mini-C plasmid. Immunofluorescence microscopy was performed to stain for endogenous cMyBP-C, and mini-C plasmid (stained with rabbit DDK tag) N = 4. (C) Co-localization was measured by Pearson’s R value, using Image J Co-Loc2 plug-in. N = 4; One-way ANOVA, Tukey’s multiple comparison test. (D) Expression of wild-type mini-C construct and missense variants within mini-C construct within myofilament prepped NRVMs was measured via immunoblotting, blotting for its DDK tag. (E) Densitometric analysis was quantified for (D) (N = 3; One-way ANOVA statistical analysis).

To investigate whether the expression of missense variants in the mini-C construct would alter myofilament binding, myofilament and supernatant protein was isolated from NRVMs transfected with wild-type Mini-C and mini-C missense variants. Mini-C was detected by its DDK tag via immunoblotting (**Fig. 2D, E**). Expression of the wild-type mini-C and mutated mini-C was detected within the myofilament protein fractions, with no significant differences between the variants. Within the supernatant samples, there was significant detection of the wild-type mini-C and the mini-C carrying the Glu1223Lys or Arg1228Cys variants.

### *MYBPHL* missense variants impair MyBP-HL sarcomere localization

To evaluate the effect of *MYBPHL* missense variants on sarcomere localization, NRVMs were transfected with wild-type MyBP-HL and MyBP-HL expressing the selected *MYBPHL* missense variants (**Fig. 3A**). Sarcomere C-zone localization was evaluated using endogenous cMyBP-C with immunofluorescence microscopy (**Fig. 3B**). All variants expressed in MyBP-HL caused decreases in sarcomere localization, with Arg285His variants causing the most significant sarcomere mislocalization (**Fig. 3C**). We also transfected neonatal rat ventricular cardiomyocytes with wild-type MyBP-HL and mutated MyBP-HL and underwent myofilament preparations. MyBP-HL protein levels within myofilament and supernatant fractions were detected with the MyBP-HL antibody via immunoblot (**Fig. 3D**). Within both myofilament and supernatant samples, we saw that missense variants did not alter MyBP-HL levels within the myofilament fraction (**Fig. 3E**).

**Figure 3:**
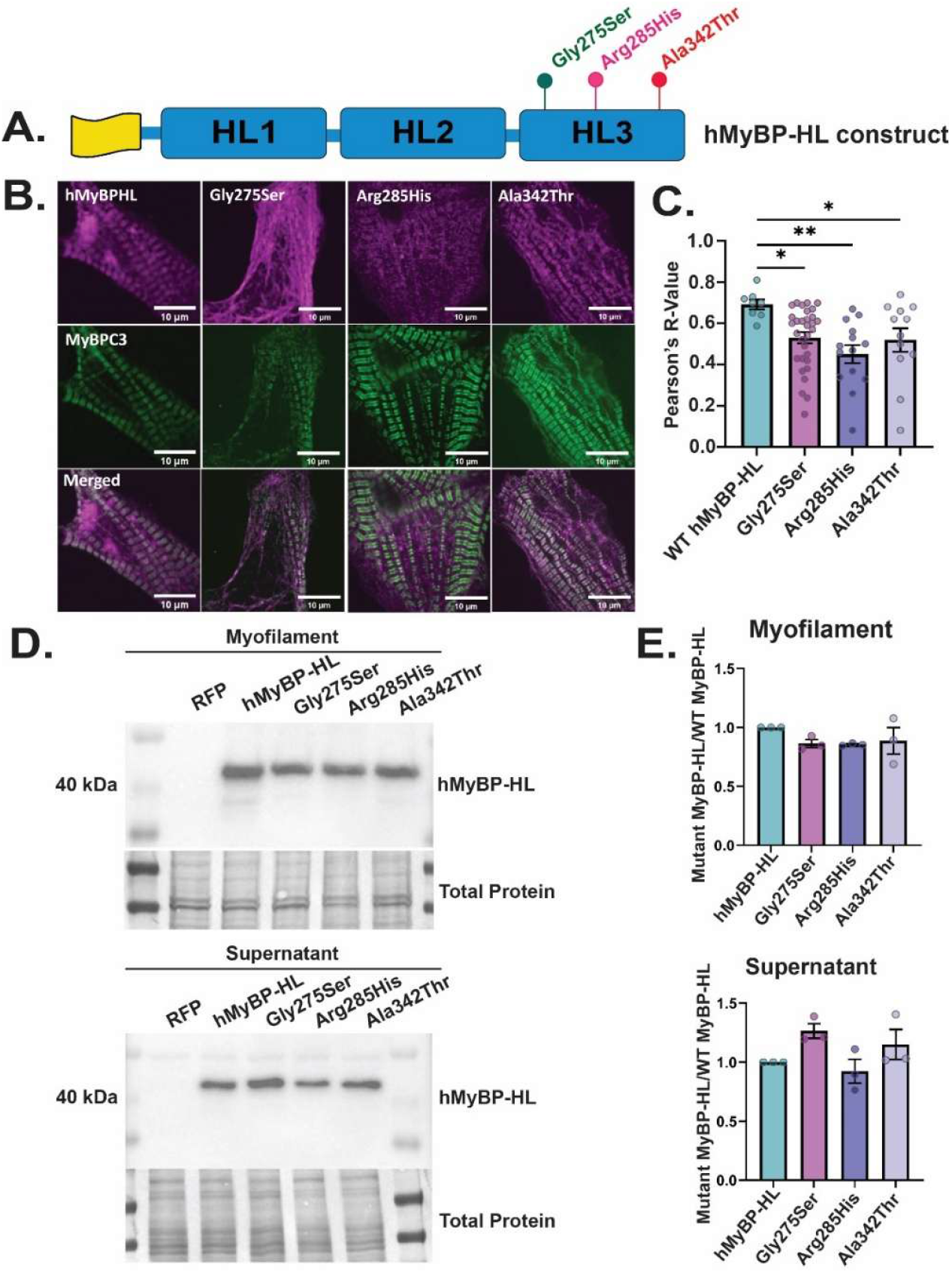
*MYBPHL* missense variants impair sarcomere localization within NRVMs. (A) Schematic showing presence of selected missense variants within MyBP-HL construct, all located in the HL3 domain. (B) NRVMs were transfected with the missense variants expressed within MyBP-HL plasmid. Immunofluorescence microscopy was performed to stain for endogenous cMyBP-C (E7), and MyBP-HL. (C) Co-localization was measured by Pearson’s R value, using Image J Co-Loc2 plug-in. N = 4; One-way ANOVA, Tukey’s multiple comparison test. (D) Expression of wild-type MyBP-HL construct and missense variants within MyBP-HL construct within myofilament prepped and supernatant samples of NRVMs was measured via immunoblotting, blotting for its DDK tag. N = 3. (E) Densitometric analysis was quantified for (D).

### A T2A/P2A system allows consistent and reproducible expression of MyBP-HL and mini-C

While mini-C and MyBP-HL constructs showed proper sarcomere localization, we needed a construct that would allow for consistent and reproducible expression of mini-C and MyBP-HL constructs at the same time to properly study the competition between myosin-binding proteins. Therefore, we created a T2A/P2A construct that contains the mini-C construct with an N-terminal DDK tag in the first position, a T2A sequence, a tdTomato fluorescent reporter in the second position, a P2A sequence, and the MyBP-HL construct in the third position (**Fig. 4A**).

**Figure 4.**
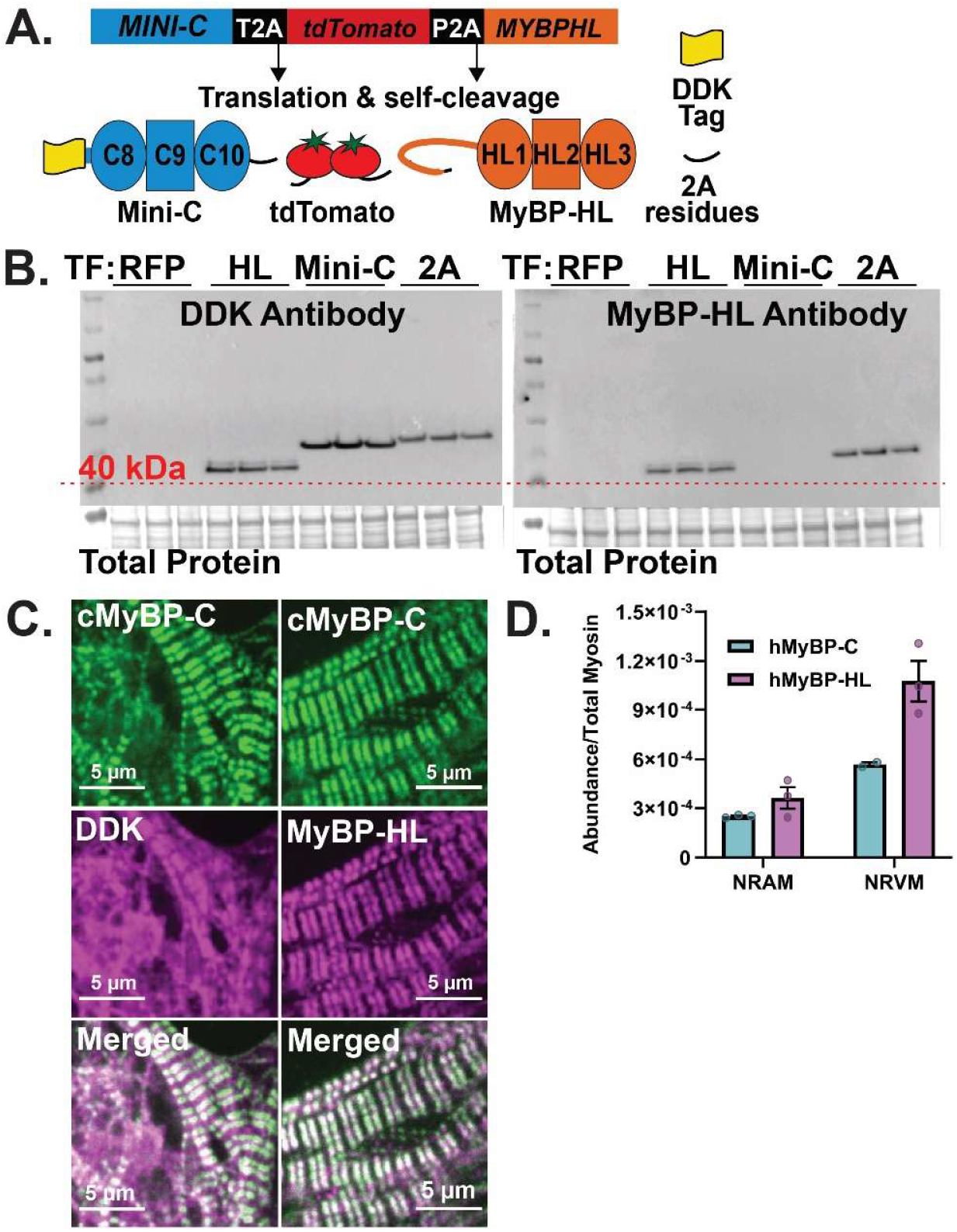
T2A-P2A construct allows for consistent and reproducible expression of both Mini-C and MyBP-HL. (A) Domain schematic showing T2A-P2A gene and its translation mechanism of producing three separate proteins, mini-C with its N-terminal DDK tag, a fluorescent reporter protein, td-tomato, and MyBP-HL. (B) Immunoblot of NRVMs expressing RFP, MyBP-HL, mini-C, and T2A/P2A constructs, showing expression of mini-C and MyBP-HL in positive controls and T2A/P2A construct (N = 3; One-way ANOVA). (C) NRVMs transfected with T2A/P2A construct, showing proper sarcomere localization of mini-C and MyBP-HL using endogenous cMyBP-C. (D) Abundances of mini-C and MyBP-HL normalized to myosin 6 and 7 in NRAMs and NRVMs transfected with T2A/P2A construct (N = 3). (E) *MYBPC3* peptide standard curve, measuring abundances of cMyBP-C within positive and negative controls, showing abundances are within range of peptide standard curve.

To validate this system, neonatal rat ventricular cardiomyocytes were transfected with RFP, serving as a negative control, plasmids with either MyBP-HL or mini-C alone as positive controls, and the T2A/P2A constructs. Using a DDK antibody, we validated expression of the independent MyBP-HL and mini-C constructs (**Fig. 4B**). Within the 2A transfected samples, the DDK tag is solely on the N’-terminal of the mini-C. The MyBP-HL antibody detected MyBP-HL in samples transfected with MyBP-HL alone, and in the T2A/P2A transfected samples, indicating expression of the MyBP-HL. Importantly, we did not identify any high molecular weight products in the T2A/P2A group, indicating that the self-cleaving 2A sites effectively generate the proteins of interest. To determine if the mini-C and MyBP-HL expressed in T2A/P2A construct would properly incorporate into the sarcomere, NRVMs were transfected with the T2A/P2A construct and immunostained for DDK to detect mini-C detection, MyBP-HL, and endogenous cMyBP-C (**Fig. 4C**). When the T2A/P2A construct was expressed in NRVMs, there was proper sarcomere incorporation of both the mini-C and MyBP-HL proteins, as seen by colocalization with endogenous cMyBP-C.

To study the competition between MyBPs in the presence of missense variants, the T2A/P2A construct needed to allow reproducible amounts of each protein. We transfected neonatal atrial and ventricular cardiomyocytes with the T2A/P2A construct and detected the abundances of mini-C and MyBP-HL with mass spectrometry (**Fig. 4D**). Abundances of mini-C and MyBP-HL peptides were normalized to the total abundance of α- and β-myosin heavy chain peptides within the samples. The normalized abundances of mini-C and MyBP-HL to myosin were consistent across NRAMs and NRVMs transfected with the T2A/P2A construct, indicating the T2A/P2A construct allows for reproducible expression of both mini-C and MyBP-HL. It is important to note that in both NRAMs and NRVMs, the ratio of MyBP-HL to myosin is higher than the ratio of mini-C to myosin. However, peptide fragments are not uniformly generated from the sample and/or detected by the spectrometer, and therefore this data cannot report the relative amount of mini-C to MyBP-HL generated from the 2A system. However, we conclude the reproducibility of the system is sufficient to model missense variants with the wild-type sequences as the control.

### *MYBPC3* missense variants in T2A/P2A show reduced sarcomere incorporation

Using the T2A/P2A construct, tested whether *MYBPC3* or *MYBPHL* missense variants could alter myosin binding protein stoichiometry. We expressed the *MYBPC3* missense variants in mini-C using the T2A/P2A construct (**Fig. 5A**). Wild-type T2A/P2A and missense mini-C T2A/P2A were transfected in NRVMs, myofilament protein fractions were prepared, and levels of mini-C and MyBP-HL within the myofilament and supernatant protein fractions were analyzed (**Fig. 5B, C**). We hypothesized that the supernatant fraction would contain missense mutant protein if it were unable to incorporate into the sarcomere. However, we found extreme variability in the supernatant fraction, even with the wild type construct. We interpret this as variability within transfection of non-cardiomyocytes. Therefore, the supernatant fraction was not quantified. To ensure that changes in MyBP protein abundance were due to the presence of missense variants and not expression levels, we immunoblotted for the tdTomato protein containing the cleaved 2A sequence for normalization. Within the myofilament fraction, mini-C missense variants Pro1181Ala/2A and Glu1223Lys/2A were reduced relative to the wild-type/2A ratio. We expected that any mutation that reduced the levels of mini-C would result in an increase in MyBP-HL. However, we found no increase in MyBP-HL/2A levels in the Pro1181Ala and Glu1223Lys samples. We expect that in this system, the amount of mini-C or MyBP-HL is not sufficient to fully outcompete endogenous cMyBP-C, and therefore we only detect the relative reduction in binding of the mutant protein and not the concomitant increase in MyBP-HL. Although there were no major changes in MyBP-HL/2A levels, we next attempted to normalize the mini-C/2A ratio to the MyBP-HL/2A ratio to account for any competition between the two proteins that was not apparent with the single protein normalization. This metric showed significant reduction in the mini-C levels in three of the *MYBPC3* mutants.

**Figure 5:**
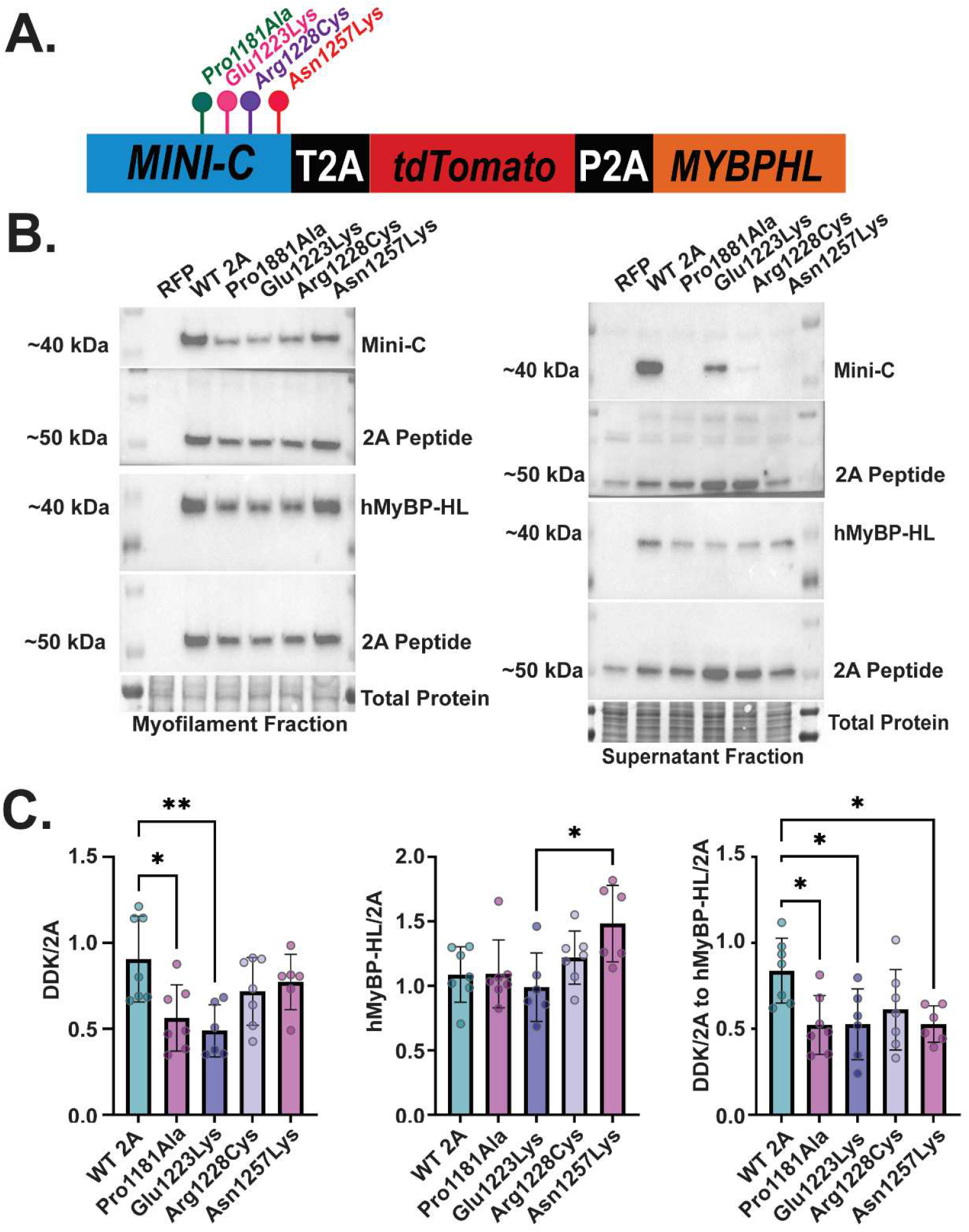
*MYBPC3* missense variants in the T2A/P2A show reduced sarcomere incorporation compared to wild type. (A) Schematic of T2A/P2A construct expressing the *MYBPC3* missense variants. (B) Representative immunoblots with myofilament and supernatant protein lysates from NRVMs transfected with the wild-type T2A/P2A construct and T2A/P2A with *MYBPC3* missense variants, blotting for mini-C, MyBP-HL, and 2A-tdTomato. (C) The ratios of DDK and MyBP-HL were normalized to their 2A expression patterns. A stoichiometry ratio between DDK/2A and MyBP-HL/2A was quantified. N = 6 – 7; One-way ANOVA, Tukey’s multiple comparison test.

### The MYBPHL Gly275Ser missense variant shows reduced myofilament incorporation

To investigate whether *MYBPHL* missense variants could change myosin binding protein stoichiometry, we transfected in NRVMs with wild-type and the three *MYBPHL* missense variants in the T2A/P2A construct (**Fig. 6A**). Protein levels of mini-C and MyBP-HL in myofilament fraction was analyzed by immunoblotting (**Fig. 6B**). To ensure that changes in myofilament abundance were not due to variable levels of construct expression we normalized the myosin binding proteins to the 2A sequence on the tdTomato (**Fig. 6C**). The Gly275Ser *MYBPHL*/2A abundance was significantly reduced compared to wild-type. Surprisingly, we saw a reduction in mini-C as well in this sample. However, normalizing the MyBP-HL/2A to the mini-C/2A showed a significant reduction, indicating that the Gly275Ser MyBP-HL was significantly less able to bind to the myofilament.

**Figure 6:**
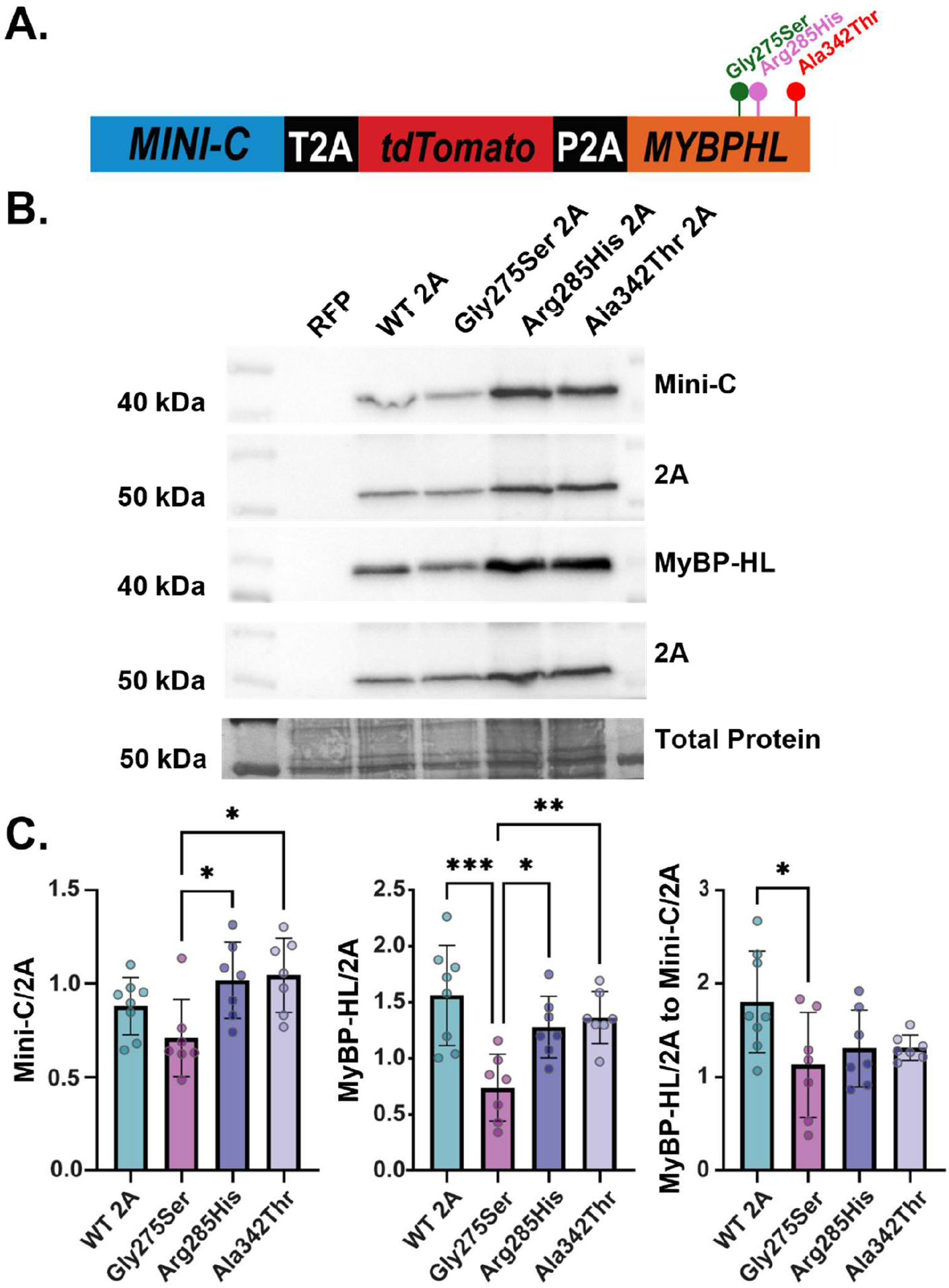
*MYBPHL* Missense Variant Gly275Ser in the T2A/P2A significantly reduces MyBP-HL sarcomere incorporation. (A) Schematic of T2A/P2A construct expressing the *MYBPHL* missense variants. (B) Representative immunoblots with myofilament protein from NRVMs transfected with the wild-type T2A/P2A construct and *MYBPHL* missense variant expressing T2A/P2A, blotting for mini-C, MyBP-HL, and the 2A peptide fragment on the tdTomato protein. (C) The ratios of DDK and MyBP-HL were normalized to the 2A protein levels. The ratio between MyBP-HL/2A and DDK/2A was quantified. N = 7 – 8; One-way ANOVA, Tukey’s multiple comparison test.

## Discussion

Mutations in sarcomere proteins have been shown to cause development of hypertrophic and dilated cardiomyopathies [8, 20]. Nonsense mutations within cMyBP-C causes early termination of translation, leading to the degradation of the mRNA via nonsense-mediated decay, preventing any functional protein from being made and causes disease by haploinsufficiency [21, 22]. In contrast, missense mutations within cMyBP-C can cause various effects, depending on the localization of the mutation within the protein [19]. Some missense mutations within the thick filament binding domain of cMyBP-C disrupt thick filament localization [22]. Due to their variability, pathogenicity of missense mutations within cMyBP-C are difficult to classify.

Furthermore, studies on *MYBPC3* missense mutations have focused on their effects in the ventricles, where cMyBP-C is the only myosin binding protein. In the context of the ventricles, if missense mutant cMyBP-C is present and has a reduced binding affinity, there are only wild-type cMyBP-C molecules that can compete with the mutant for myofilament incorporation. This leads to a relatively normal phenotype as one allele of cMyBP-C is nearly sufficient to maintain normal sarcomere function [23, 24]. However, in the atria, cMyBP-C and MyBP-HL compete for sarcomere binding sites and maintain a roughly equal ratio [17]. Missense mutations in cMyBP-C that reduce thick filament binding would compete with the wild-type allele of cMyBP-C and two alleles of wild-type MyBP-HL. This could cause a shift in myosin binding protein stoichiometry and potentially alter atrial contractility.

Our results suggest that certain *MYBPC3* and *MYBPHL* missense variants in the thick filament binding domains can cause mislocalization from the C-zone of the sarcomere while still maintaining some myofilament binding. The sarcomere is a highly structured system, and its function is not solely dictated by the presence of specific sarcomere proteins but by their proper localization and stoichiometric balance [25-27]. We show that missense variants in myosin binding proteins reduce their affinity for their native binding sites. This is particularly important within the context of the atria, where cMyBP-C and MyBP-HL are co-expressed, and the mislocalization of one protein can alter their stoichiometric equilibrium, potentially disturbing the finely tuned contractile mechanisms.

Thus, our findings suggest that myosin binding protein missense variants that cause mislocalization with aberrant sarcomere binding could represent an underappreciated mechanism for early or subclinical stages of cardiomyopathy, especially regarding atrial dysfunction. Our work highlights the need for studies of atrial biophysics and physiology in the context of myosin binding protein-linked cardiomyopathies.

## Abbreviations

(DCM): Dilated Cardiomyopathy
(HCM): Hypertrophic Cardiomyopathy
(AF): Atrial Fibrillation
(MyBP-HL): Myosin-binding protein-H like
(cMyBP-C): Cardiac myosin binding protein-C
(NRVMs): Neonatal rat ventricular cardiomyocytes
(NRAMs): Neonatal rat atrial cardiomyocytes

## Limitations

While this study provides insights on the effects of missense variants on myosin binding protein stoichiometry, we are limited by the inability to analyze the expression of MyBPs as a ratio of myofilament to supernatant due to the heterogeneity of the cell population in our transfection approach. When cardiomyocytes are isolated for transfection and immunoblotting experiments, there are inevitably some percentage of fibroblasts, endothelial cells, and smooth muscle cells even when steps are taken to improve the purity of the cardiomyocytes. This, in addition to the low and variable transfection efficiency of neonatal rat cardiomyocytes makes it impossible to evaluate how much myosin binding protein is myofilament-bound compared to how much is soluble. Future experiments could include expressing the T2A/P2A construct via viral delivery in a cardiomyocyte lacking endogenous myosin binding proteins.

## Author contributions

K.N.A. and D.Y.B. conceived and designed research; K.N.A., H.E.C., Y.Y., G.E.F., T.B., A.P., and L.M.W. performed experiments; K.N.A., H.E.C., and D.Y.B. analyzed data; K.N.A. and D.Y.B. interpreted results of experiments; K.N.A., H.E.C., and D.Y.B. prepared figures; K.N.A. and D.Y.B. drafted the manuscript; K.N.A. and D.Y.B. edited and revised the manuscript; all authors approved the final version of the manuscript.

